# Atrioventricular nodal reentrant tachycardia onset, sustainability, and spontaneous termination in rabbit atrioventricular node model with autonomic nervous system control

**DOI:** 10.1101/2024.06.10.598392

**Authors:** Maxim Ryzhii, Elena Ryzhii

## Abstract

Atrioventricular nodal reentrant tachycardia (AVNRT) is one of the most common types of paroxysmal supraventricular tachycardia. The autonomic nervous system (ANS) activity is known to affect sudden episodes of AVNRT, but the detailed underlying mechanism is not fully understood. In this work, we update our recent compact multifunctional model of the rabbit atrioventricular node (AV) with ANS control to simulate AVNRT. The refractoriness of model cells is modulated by one ANS coefficient, causing a change in the effective refractory periods, conduction delays, and intrinsic frequency of pacemaker cells. Using the model, we examine the onset, sustainability, and spontaneous termination of typical slow-fast and atypical fast-slow forms of AVNRT under ANS modulation. The conditions for the onset and sustainability of AVNRT can occur independently of each other in various combinations. Differences in the effective refractory periods of the slow and fast pathways of the AV node during anterograde and retrograde conduction determine the form of AVNRT. For the first time, the possibility of identifying hidden processes occurring inside the AV node using a computer model is shown, allowing us to come closer to understanding the role of ANS control during AVNRT. The results obtained are consistent with clinical and experimental data and represent a new tool for studying the electrophysiological mechanisms of this type of arrhythmia.

## 1 INTRODUCTION

The atrioventricular (AV) node consists of dual pathways: a fast pathway (FP) with a relatively longer effective refractory period (ERP) and a slow pathway (SP) with a shorter ERP. These pathways can create a reentrant circuit, a substrate for AV nodal reentrant tachycardia (AVNRT), the most common type among regular supraventricular arrhythmias (29).

AVNRT manifests as sudden episodes of a few cycles of abnormally fast heartbeats (reciprocal or echo beats) or as sustained or persistent tachycardia. AVNRT is electrophysiologically classified as typical (slow-fast) and atypical (fast-slow, and also slow-slow in humans) forms corresponding to anterograde-retrograde conduction sequence through the dual AV nodal pathways (15).

The autonomic nervous system (ANS) plays an essential role in initiating and terminating supraventricular tachycardias in the AV node (20). Sympathetic stimulation usually facilitates the induction of AVNRT, whereas enhancement of vagal (parasympathetic) tone through pharmacological administration or Valsalva maneuvers is commonly used to terminate the tachycardia (2, 34). The effect of ANS control on dual pathways interaction in the initiation, sustainability, and spontaneous termination of AVNRT is still poorly understood despite some attempts to explain its exact underlying physiological mechanism.

A few functional computer models of AV node were developed (13, 7, 21), but to our knowledge, only one of them incorporates ANS control (21). In the latter model, the authors modulate vagal tone by modifying parameters of AV node refractoriness and conduction velocity separately. Recently, we have developed a compact, multi-functional rabbit AV node model based on the simplified two-variable cardiac cell model (25). The one-dimensional model includes dual pathways, primary pacemaker in the sinoatrial node (SN), and subsidiary pacemaker in SP and is tuned to the available experimental data. Visualization of interactions between intact and post-ablated SP and FP in the form of Lewis ladder diagrams facilitates the study of AVNRT.

Experimental observations show that FP has a significantly longer effective refractory period (ERP) in the case of anterograde conduction (aERP*_F P_*) than that of SP (aERP*_SP_*), which is a substrate for typical slow-fast AVNRT at stimulation periods shorter than the aERP*_F P_* (22). This substantial property of normal AV node behavior, demonstrated by simulation and experimental studies (13, 7, 3), provides effective conduction slowing and fast rhythm filtering. In contrast, in the case of retrograde conduction with His bundle pacing no apparent difference between retrograde ERPs of SP (rERP*_SP_*) and FP (rERP*_F P_*) was observed in both control and post-ablation cases. Taking into account the statistical uncertainty of the difference between rERP*_SP_* and rERP*_F P_* (22), in our preliminary work (26), we assumed that any relationship between rERP*_SP_* and rERP*_F P_* values may exist within a reasonable range. We showed that rERP*_SP_* and rERP*_F P_* affected insignificantly the anterograde conduction in the AV node. Along with the typical AVNRT of atrial origin, we simulated both typical and atypical AVNRT forms with His bundle pacing and demonstrated that the difference in aERP and rERP of FP and SP determines the form of AVNRT. However, these results did not take into consideration the influence of ANS control.

To overcome this drawback, we updated the functionality of our AV node model incorporating the ANS control. The sympathetic and parasympathetic control was implemented by including only one coefficient for scaling parameters related to the refractoriness of model cells. Other main cardiac conduction system properties, such as intrinsic rates of pacemakers and conduction times, depend on this coefficient.

In the current work, using the modified model, we study the onset, susceptibility, and spontaneous termination of typical slow-fast and atypical fast-slow forms of AVNRT. We consider the induction of AVNRT not only by premature atrial and His bundle stimulation, referred to in clinical practice as premature atrial (PAC) and ventricular (PVC) complexes, but also by electrical impulses originating within AN node (premature junctional complex, PJC) (29).

## MODEL AND METHODS

The scheme of the compact AV node model used in this study is shown in Fig. 1(A). Each model cell is described by Aliev-Panfilov cardiac cell model (1) given by a couple of reaction-diffusion type ordinary differential equations

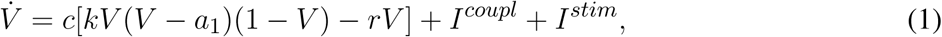

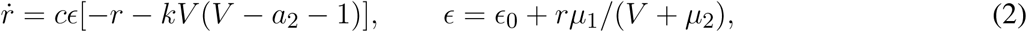

**Figure 1.**
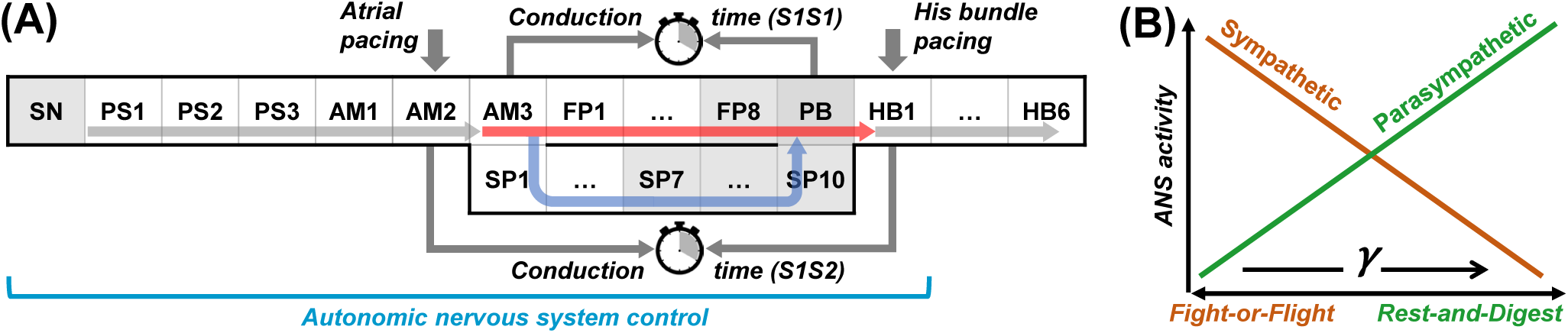
(A) Schematic representation of the rabbit atrioventricular node model. SN - sinus node, PS - peripheral sinus node cells, AM - atrial muscle cells, FP - fast pathway cells, SP - slow pathway cells, PB - penetrating bundle cell, HB - His bundle cells. Thick vertical arrows denote places of stimuli application for atrial and His bundle pacing. The arrows within the structure correspond to normal conduction. The gray shading indicates pacemaker cells. (B) Relation between ANS activity and the coefficient *γ*.

where *V* is the dimensionless transmembrane potential, *r* is the gate variable, *c* is the time scaling coefficient, and *k* is the parameter controlling the magnitude of the transmembrane current. Parameters *ɛ*_0_, *a*_1_, *a*_2_, *µ*_1_, and *µ*_2_ determine the conduction characteristics of tissue, *a*_1_ *>* 0 represent the excitation threshold of quiescent excitable cells, while *a*_1_ *<* 0 sets the intrinsic oscillation frequency of the pacemaking cells [gray-shaded in Fig. 1(A)] (24). The intercellular coupling terms

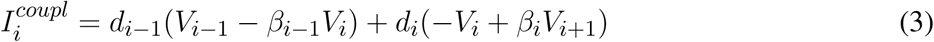

in the one-dimensional system account for the coupling asymmetry, where *d_i_* are the diffusion coefficients (normalized on dimensionless distance), *i* denotes the index of neighboring cells, and the coefficients *β <* 1 correspond to the accelerated anterograde and slowed retrograde conduction, and vice versa for *β >* 1. *I^stim^* denotes external stimulation current applied to the atria or His bundle [thick arrows in Fig. 1(A)] to perform S1S2 and S1S1 stimulation protocols.

Standard S1S2 stimulation consisted of nine pulses with constant basic S1–S1 interval equal to spontaneous sinus rhythm interval determined by the current ANS state, and S2 test premature stimulus with S1–S2 interval introduced with a decrement of 1 ms until conduction through AV node is blocked. The S1S2 conduction time was measured between AM2 and HB1 model cells. S1S1 stimulation was performed by applying ten pulses with 1 ms interval decrement starting from from the interval of spontaneous sinus rhythm in the current state of ANS. For this stimulation type, we measured atria-His and His-atria conduction delays within the AV node ring between AM3 and PB model cells. Atrial and His bundle stimulation pulses were 1 ms and 2 ms long, respectively, and 1.3 times above the threshold.

We implemented the effect of the ANS in our rabbit cardiac conduction system model by the introduction of a control coefficient *γ* [Fig. 1(B)], allowing dynamical changing of the parameters *µ*_1_ and *µ*_2_ responsible for the cell refractoriness (1):

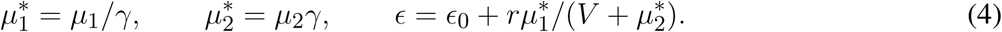

The rationale for this method of ANS control was discussed in detail in our recent report (27). To reflect different states of the ANS, the same coefficient *γ* was simultaneously applied in all model cells of the part of the cardiac conduction system [Fig. 1(A)] from the sinus node (SN) to the penetrating bundle (PB), including the AV junctional pacemaker (10).

Since only isolated rabbit heart preparations were used in the experiments (22, 7, 3), any influence of the ANS was absent, leaving the hearts in a state representing a static invariable situation regarding the cardiac conduction system. At such conditions, the onset of AVNRT was observed in the S1S2 protocol stimulation at short atrial test pulses with a normal sinus rhythm of about 166 bpm (360 ms beating interval) (22). However, it is known that enhancing sympathetic tone may provoke AVNRT onset (12). At the same time, Valsalva maneuver or adenosine administration (2, 34) causes the vagal tone enhancement, resulting in reduced heart rate and consequent termination of AVNRT. Considering the above facts, we set *γ* = 1.0 corresponding to an augmented sympathetic tone state with high sinus rhythm, allowing induction of some form of AVNRT. An increase of *γ* first leads to a normal rhythm at *γ* ≃ 1.7 and then to bradycardia when the vagal tone strongly predominates (*γ* ≃ 2.0). A decrease of *γ* means enhancement of sympathetic tone and, respectively, shortens the refractory period, action potential duration of the affected model cells, and ERP of both pathways, reduces nodal conduction time, and increases intrinsic rates of the sinus node and AV nodal pacemakers (6).

The three AV node model variants considered had similar anterograde conduction characteristics (aERP*_SP_ <* aERP*_F P_*) but different retrograde conduction properties (26):

– the *first model variant* had rERP*_SP_* ≃ rERP*_F P_* for enhanced sympathetic tone (increased sinus rhythms) and at normal condition;
– the *second model variant* had slightly reduced diffusion coefficient *d* for the last three FP cells and increased *d* for the last four SP cells compared with the first variant which provides longer rERP*_F P_* within the entire *γ* range;
– the *third model variant* had the coupling asymmetry coefficient *β* reduced five times between SP10 and PB model cells, creating the relationship rERP*_SP_ >* rERP*_F P_* at enhanced sympathetic tone (higher rhythms) and its inversion (rERP*_SP_ <* rERP*_F P_*) at normal condition and enhanced vagal tone (reduced rhythms). The refractory period in the proximal part of His bundle (HB1–HB3 cells) was reduced in the second and third model variants compared to the first variant to facilitate His bundle premature stimulation.

For each model variant, we considered three scenarios of premature cardiac complexes classified by their origin - atrial (premature atrial complex, PAC), ventricular or His bundle (premature ventricular complex, PVC), and AV intranodal or junctional (premature junctional complex, PJC). The cases of PAC and PVC required proper selection of intervals of premature extrastimulus within the AVNRT induction window. For PJC, a short burst of sympathetic activity with high sinus tachycardia was applied to stimulate the conduction block within the AV node.

The ANS coefficient *γ* was dynamically varied stepwise during the simulations at predefined moments. The intervals between *γ* changes were selected to reflect the natural reaction of the cardiac conduction to ANS modulation.

The simulations were conducted with MATLAB (R2023a, Mathworks Inc., Natick, MA, USA). The ordinary differential equations were solved using ode23 solver which utilizes second and third order Runge-Kutta-Fehlberg formulas with automatic step-size. Other parameter values were similar to that in (25) and are based on rabbit experimental data (22, 3). Additional details of the basic rabbit AV node model and its properties can be also found in (25).

## 2 RESULTS

In what follows, we refer to AVNRT as sustained oscillations within the AV ring and echo beats (reciprocating pulses) as decaying oscillations with with no more than a few cycles.

### 2.1 The First Model Variant

Figure 2 demonstrates simulation results of AVNRT onset and spontaneous termination for the first model variant with varying ANS tone (*γ*). The top panels demonstrate ladder diagrams of conduction propagation from the sinus node to His bundle via FP (red) and SP (blue color). The value of the current ANS coefficient *γ* and the corresponding sinus rate in beats per minute (in brackets) are indicated above the ladder diagrams. Vertical dashed lines denote the moments of *γ* change. The middle and bottom panels show action potential sequences from the sinus node to His bundle separately for FP and SP.

**Figure 2.**
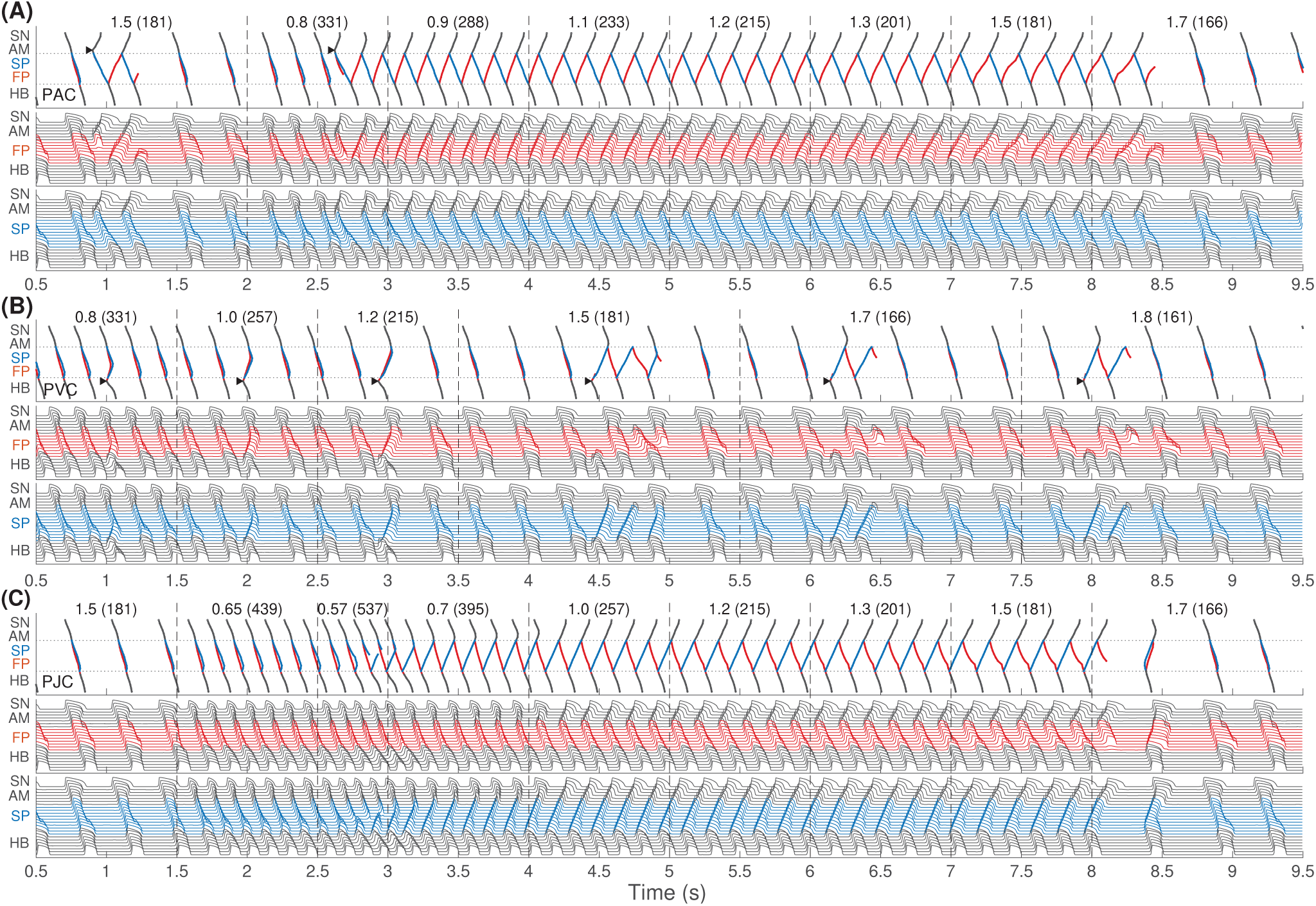
The first model variant with rERP*_SP_* ≃ rERP*_F P_*. The top panels show the ladder diagrams of excitation propagation in the AV node dual pathway structure from the sinus node (SN) to the His bundle (HB) via fast pathway (FP, red traces) and slow pathway (SP, blue traces). The numbers above the diagrams indicate the value of the coefficient *γ* and the corresponding sinus rhythm in bpm (in brackets). Black arrowheads denote the places and moments of premature stimulation. The middle and bottom panels demonstrate action potentials passing through the fast and slow pathways. (A) Onset of typical slow-fast AVNRT with premature atrial stimulation (premature atrial complex, PAC) at enhanced sympathetic tone followed by spontaneous termination at rising vagal tone. (B) No induction of AVNRT was observed with His bundle pacing (premature ventricular complex, PVC). (C) A brief burst of very strong sympathetic tone accompanied by an excessively accelerated sinus rhythm (premature junctional complex, PJC) induced atypical fast-slow form of AVNRT followed by spontaneous termination with vagal tone dominance.

In Fig. 2(A), we started with an increased sympathetic tone at *γ* = 1.5 applying premature atrial stimulus (PAC, indicated by black arrowhead), which caused a slow-fast echo beat followed by sinus rhythm. Applying PAC at *γ* decreased to 0.8, we obtained the onset of typical slow-fast AVNRT. The oscillations persisted with increasing *γ* up to 1.5 and spontaneously terminated at *γ* = 1.7 with a return to the normal sinus rhythm. Details of slow-fast AVNRT onset with PAC are shown on the ladder diagram in Fig. 3(A).

**Figure 3.**
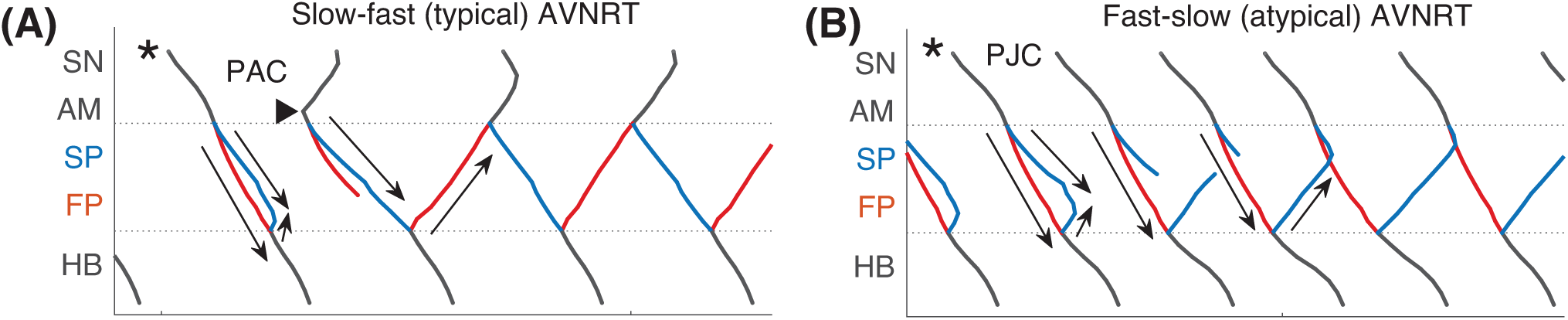
Schemes of AVNRT onset in the first model variant. (A) Slow-fast (typical) AVNRT form at the premature atrial complex (PAC) [from Fig. 2(A)], and (B) fast-slow (atypical) form at the premature junctional complex (PJC) [from Fig. 2(C)]. Arrows show the direction of conduction via pathways. Arrowhead denotes the place and moment of premature stimulation. Asterisk indicates normal conduction.

With His bundle premature stimulation (PVC), trying to provoke AVNRT at different *γ*, we obtained only atypical fast-slow echo beats during at *γ* ≥ 1.5 [Fig. 2(B)].

AVNRT with PJC originates from within the AV node, so it does not require an external premature stimulus. A brief episode with a sudden decrease of *γ* to the very low value of 0.57, accompanied by a very high sinus rhythm of 537 bpm, resulted in a critical reduction in the duration and amplitude of action potential and block of SP conduction. This triggered fast-slow AVNRT, which persisted with increasing *γ* until the latter reached a normal value of 1.7 [Fig. 2(C)]. Details of fast-slow AVNRT onset of PJC origin are shown on the ladder diagram in Fig. 3(B). The AVNRT began after retrograde SP excitation met anterograde SP excitation from a subsequent sinus rhythm, and they annihilated each other.

To investigate the underlying physiological background of the results in Fig. 2, we performed simulations using S1S2 and S1S1 stimulation protocols. Figure 4 presents various related conduction curves for the first model variant calculated for different ANS states represented by the coefficient *γ* = 0.8, 1.0, 1.2, 1.5, and 1.7. Figures 4(A) and 4(B) demonstrate anterograde conduction curves with PAC and retrograde conduction curves with PVC for the control (intact) AV node. With decreasing *γ*, i.e., increasing sympathetic tone, the anterograde conduction switching from FP to SP (22, 7) became more pronounced with sharper tilt [Fig. 4(A)].

**Figure 4.**
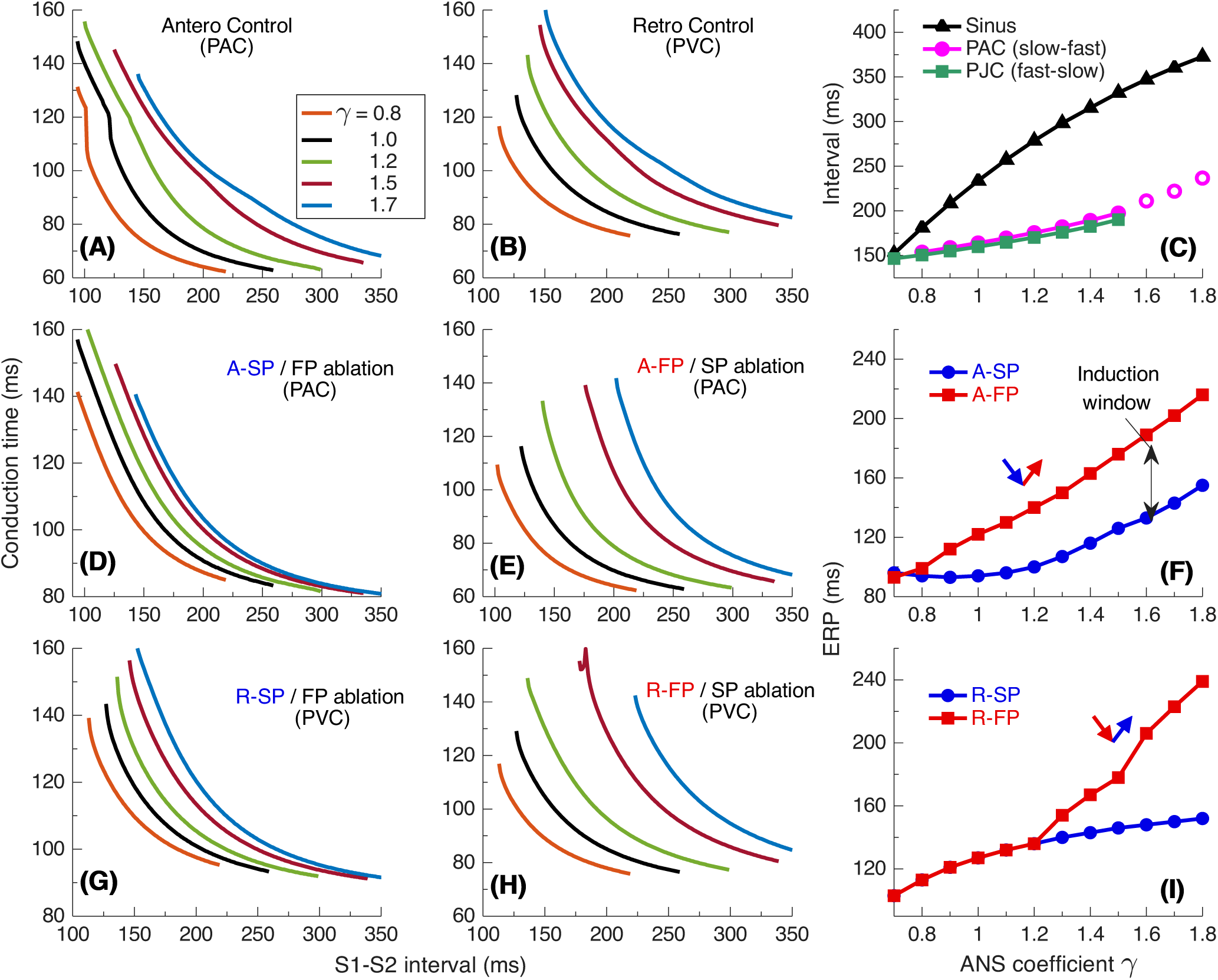
Conduction characteristics using S1S2 stimulation protocol for the first model variant. (A) Control case with atrial pacing (PAC). (B) Control case with His bundle pacing (PVC). (C) Dependence of sinus rhythm and AVNRT intervals of PAC and AV intranodal (PJC) origin on the coefficient *γ*. Empty markers correspond to echo beats. (D) and (E) - anterograde (A-) conduction times for SP (FP ablation) and FP (SP ablation), and their ERPs (F); (G) and (H) - retrograde (R-) conduction times for SP and FP, and their ERPs (I).

Maximal S1-S2 interval values for the conduction curves are limited by spontaneous sinus rhythm interval, which decreases with smaller *γ* [Fig. 4(C)]. Apart from the sinus rhythm interval, ANVRT intervals obtained with PAC and PJC shown in Figs. 2(A) and 2(C) are indicated in Fig. 4(C) by filled markers, and echo beats appearing at *γ* = 1.6 − 1.8 - by open markers. The intervals of AVNRT of different origin are always shorter than the sinus rhythm interval and indicate the overdrive suppression of the latter by the reentrant oscillations.

The S1S2 conduction curves obtained for individual SP and FP pathways (post-ablation cases) and the dependence of their ERPs on *γ* for PAC and PVC are shown in Figs. 4(D)–4(F) and 4(G)–4(I), respectively. The distance between FP-only curves [Figs. 4(E) and 4(H)] with changing *γ* is more pronounced than in the case of SP-only curves [Figs. 4(D) and 4(G)] for both atrial and His bundle stimulation. Within the entire useful range of *γ*, aERP*_SP_* is smaller than aERP*_F P_* and the incuction window (distance between FP and SP ERP curves) widens with a predominance of parasympathetic tone [Fig. 4(F)]. This is common for mammalian AV node (22) and creates a possibility of slow-fast AVNRT onset within the wide range of ANS states. On the other hand, the retrograde ERPs of both pathways (rERP*_SP_* and rERP*_F P_*) are equal within the entire range of sympathetic tone, and rERP*_F P_ >* rERP*_SP_* at enhanced parasympathetic tone. [Fig. 4(I)]. The above relationships between the SP and FP ERPs allowed the onset of slow-fast AVNRT with atrial pacing at *γ* ≥ 0.8 [Fig. 2(A)], and blocked the initiation of fast-slow AVNRT with His bundle pacing at *γ <* 1.3. However, the onset of fast-slow AVNRT or echo beats is possible at *γ >* 1.3 [Fig. 2(B)].

Figures 5(A)–5(C) show the conduction time calculated using S1S1 pacing protocol for anterograde SP, retrograde FP and their sum for slow-fast AVNRT form, and Figs. 5(D)–5(F) - anterograde FP and retrograde SP and their sum for fast-slow form.

**Figure 5.**
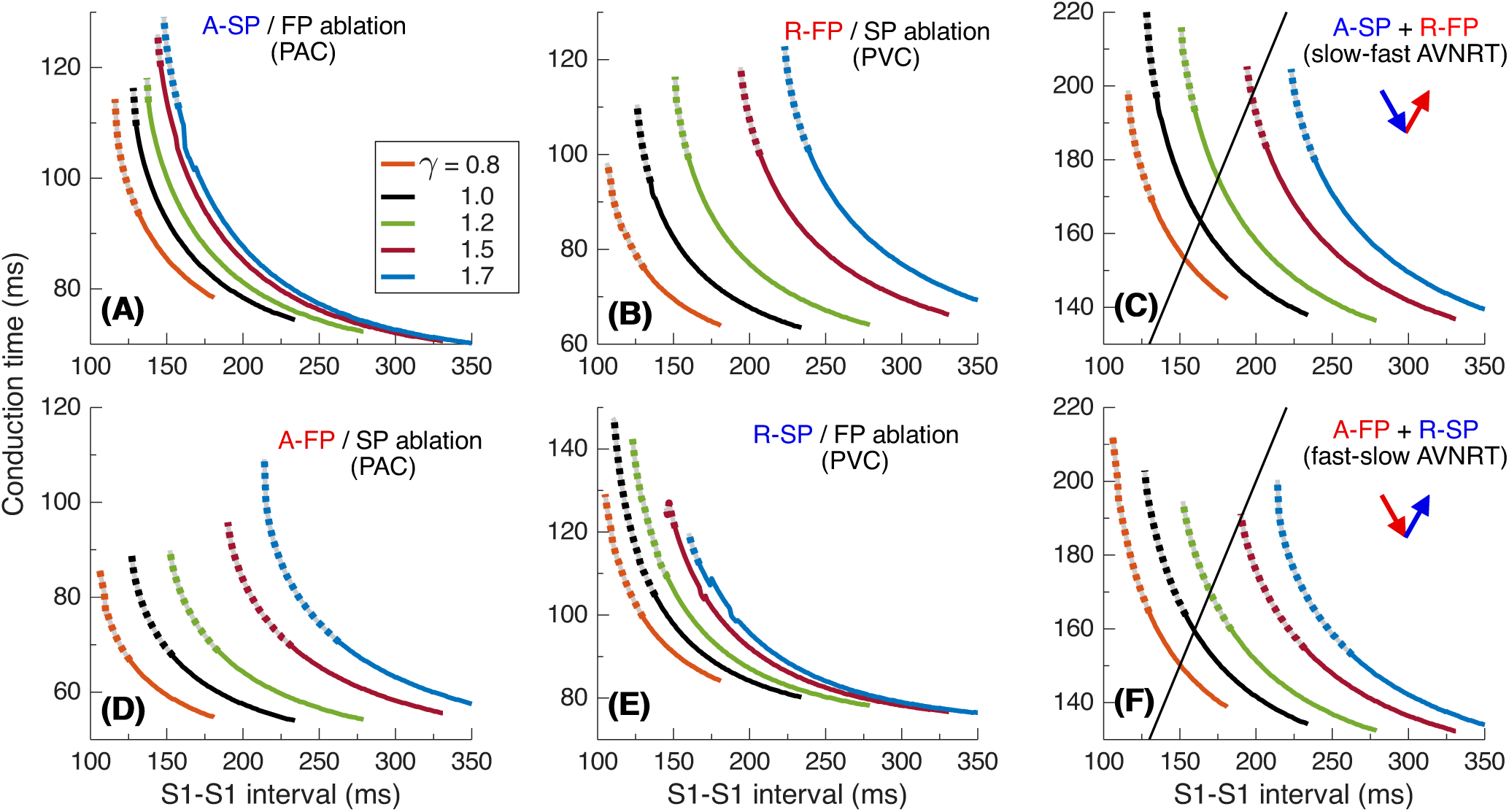
Conduction characteristics using S1S1 stimulation protocol for the first model variant. (A) and (B) - anterograde SP and retrograde FP conduction times and their sum (C) versus pacing interval. (D) and (E) - anterograde FP and retrograde SP conduction times and their sum (F) versus pacing interval. Unstable conduction is indicated by thick dotted parts on the conduction curves. Black straight lines in panels (C) and (F) represent the AVNRT sustainability lines on which the stimulation interval equals the sum of the SP and FP conduction times.

In our simulations, the initial time delay preceding the first S1 stimulus of the S1S1 protocol affected the stability of conduction in the pathways. The variation of the initial delay resulted in the unstable conduction of last S1 test stimulus at short S1-S1 intervals [left ends of the conduction curves in Figs. 5(A), 5(B), 5(D), and 5(E)]. The instability is also reflected in the summation curves in Figs. 5(C) and 5(F). The unstable regions are marked by thick dotted lines of the same color. The earliest (leftmost) point on each summation curve and the width of its unstable region are determined by the largest first unstable point and the largest last unstable point of corresponding conduction curves of either pathway.

AVNRT is a self-sustained oscillation with a cycle length equal to the stimulation period. The AVNRT cycle length equals the sum of the anterograde SP (A-SP) and retrograde FP (R-FP) conduction times for slow-fast AVNRT form and the sum of the anterograde FP (A-FP) and retrograde SP (R-SP) conduction times for fast-slow form [Figs. 5(C) and 5(F)]. At the same time, the pathway conduction time is cycle-length dependent. The existence of stable periodic oscillations in the AV ring can be determined by the presence of the intersection point of the summation curve with the identity line *y* = *x* (where S1-S1 pacing interval equals the sum of SP and FP conduction times), denoted by straight black solid lines (AVNRT sustainability lines) in Figs. 5(C) and 5(F).

As seen from Figs. 5(C) and 5(F), at *γ* = 0.8 − 1.5 the intersection points between the corresponding summation conduction curves and the AVNRT sustainability lines exist. This indicates the existence of oscillations which may be unstable at *γ* = 1.5 for slow-fast AVNRT type [see also Fig. 2(A)], and at *γ* = 1.2 − 1.5 for fast-slow type [Figs. 2(B) and 2(C)]. At *γ* = 1.7 the oscillations cannot persist for both slow-fast and fast-slow AVNRT types.

### 2.2 The Second Model Variant

Figures 6–9 present results with a similar simulation setup but for the second model variant with rERP*_SP_ <* rERP*_F P_*in the entire range of *γ*.

**Figure 6.**
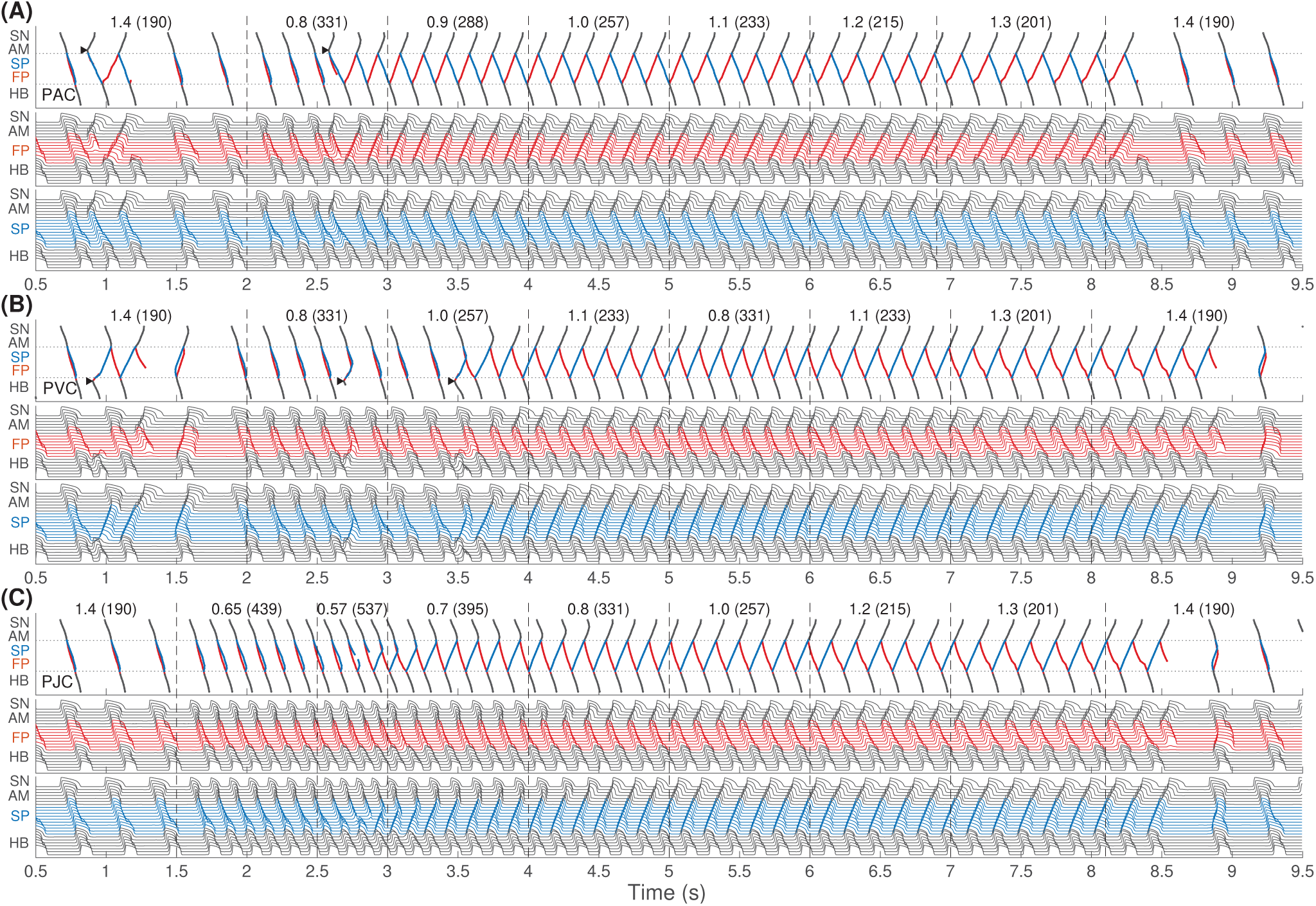
The same as in Fig. 2 but for the second model variant with rERP*_SP_ <* rERP*_F P_*. (A) Onset of slow-fast AVNRT with PAC at the enhanced sympathetic tone. At *γ* = 1.4, only slow-fast echo beats appeared. (B) Onset of the fast-slow form of AVNRT with PVC at the enhanced sympathetic tone. At *γ* = 1.4, only fast-slow echo beats appeared. (C) Similar to Fig. 2, a brief burst of very strong sympathetic tone (equivalent to PJC) induced fast-slow form of AVNRT. In all cases (A)-(C), spontaneous termination of the oscillations took place at increasing vagal tone (*γ* ≥ 1.4).

In Fig. 6(A) with atrial pacing we observed a similar situation as in Fig. 2(A), but spontaneous termination of slow-fast AVNRT occurred earlier at lower *γ* = 1.4. However, His bundle pacing gave different results [Fig. 6(B)]. First, a few fast-slow echo beats were initiated with *γ* = 1.4. Then, increasing *γ* from low values, we managed to induce fast-slow AVNRT only at *γ* ≥ 1.0. Sustained oscillations in the AV ring continued until *γ* reached 1.4. It was also possible to induce a fast-slow form of AVNRT with PJC by briefly decreasing *γ* to 0.57 in the same way as for the first model variant. In this case, the oscillations persisted with increasing *γ* up to 1.4 [Fig. 6(C)].

Figure 7 demonstrates on the ladder diagram the details of the onset of fast-slow AVNRT with His bundle pacing (PVC). The beginning of the oscillations was facilitated by a subsequent sinus impulse, similar to the situation with PJC-originated fast-slow AVNRT shown in Fig. 3(B).

**Figure 7.**
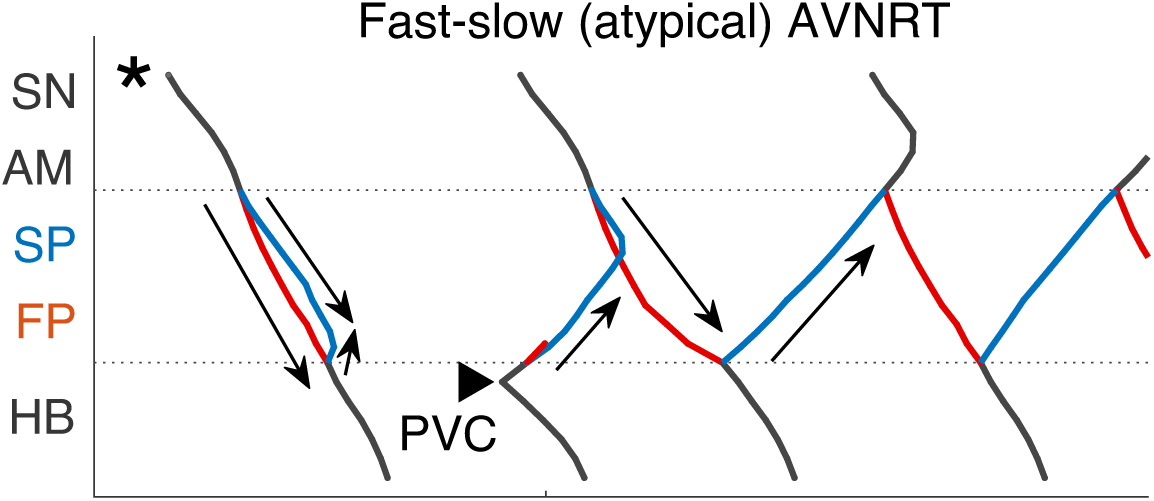
Scheme of the fast-slow AVNRT onset with PVC in the second model variant [from Fig. 6(B)]. Arrows show the direction of conduction via pathways. Arrowhead denotes the place and moment of premature stimulation. Asterisk indicates normal conduction.

The anterograde S1S2 conduction curves in Figs. 8(A), 8(D), and 8(E) and the relationship of anterograde ERPs between SP and FP [Fig. 8(F)] remained similar to those of the first model variant [Fig. 4(A), 4(D), 4(E), and 4(F)]. However, in contrast to Fig. 4(B), noticeable transitions of the conduction from FP to SP appeared on the retrograde control curves [Fig. 8(B)]. They became sharper with decreasing *γ*, due to the increase of rERP*_F P_* introduced in this model variant and reflected in Fig. 8(I) with relatively wide induction window for fast-slow AVNRT.

**Figure 8.**
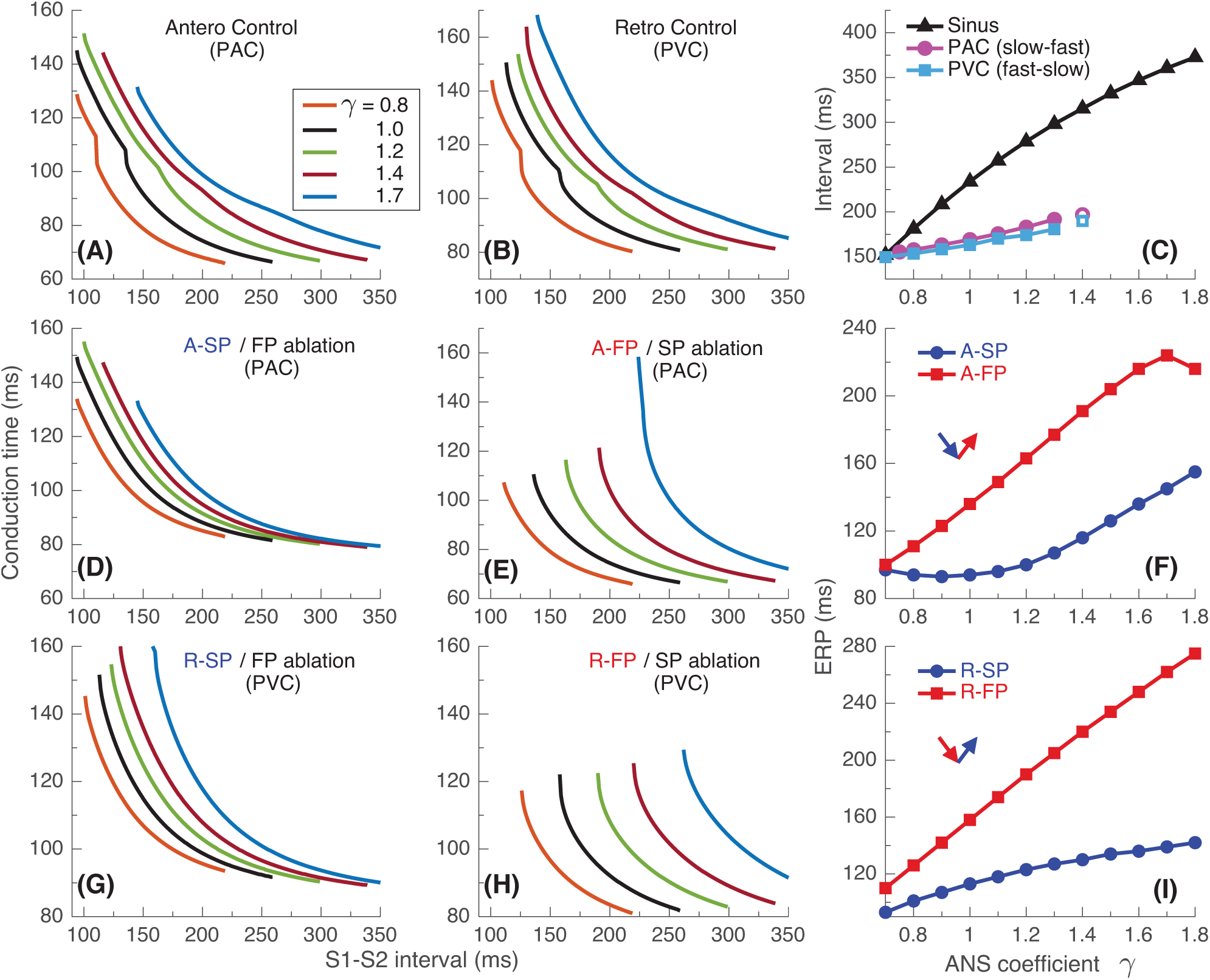
Conduction characteristics using S1S2 stimulation protocol for the second model variant.

Figure 9 shows the conduction characteristics using S1S1 stimulation protocol similar to that shown in Fig. 5. As seen from Figs. 6(A) and 9(C), the sustainability of slow-fast AVNRT persisted only at *γ <* 1.4, that is, in the narrower range of *γ* than in the case of first model variant [Fig. 5(C)]. In the cases of His bundle pacing [PVC, Fig. 6(B)] and PJC [Fig. 6(C)] the upper sustainability limit for fast-slow AVNRT case was also *γ <* 1.4 [Fig. 9(F)]. It should be noted that with His bundle pacing at *γ* = 0.8, neither AVNRT nor echo beats were induced [Fig. 6(B)], while fast-slow AVNRT initiated at higher *γ* values persisted when *γ* was temporarily reduced to 0.8.

**Figure 9.**
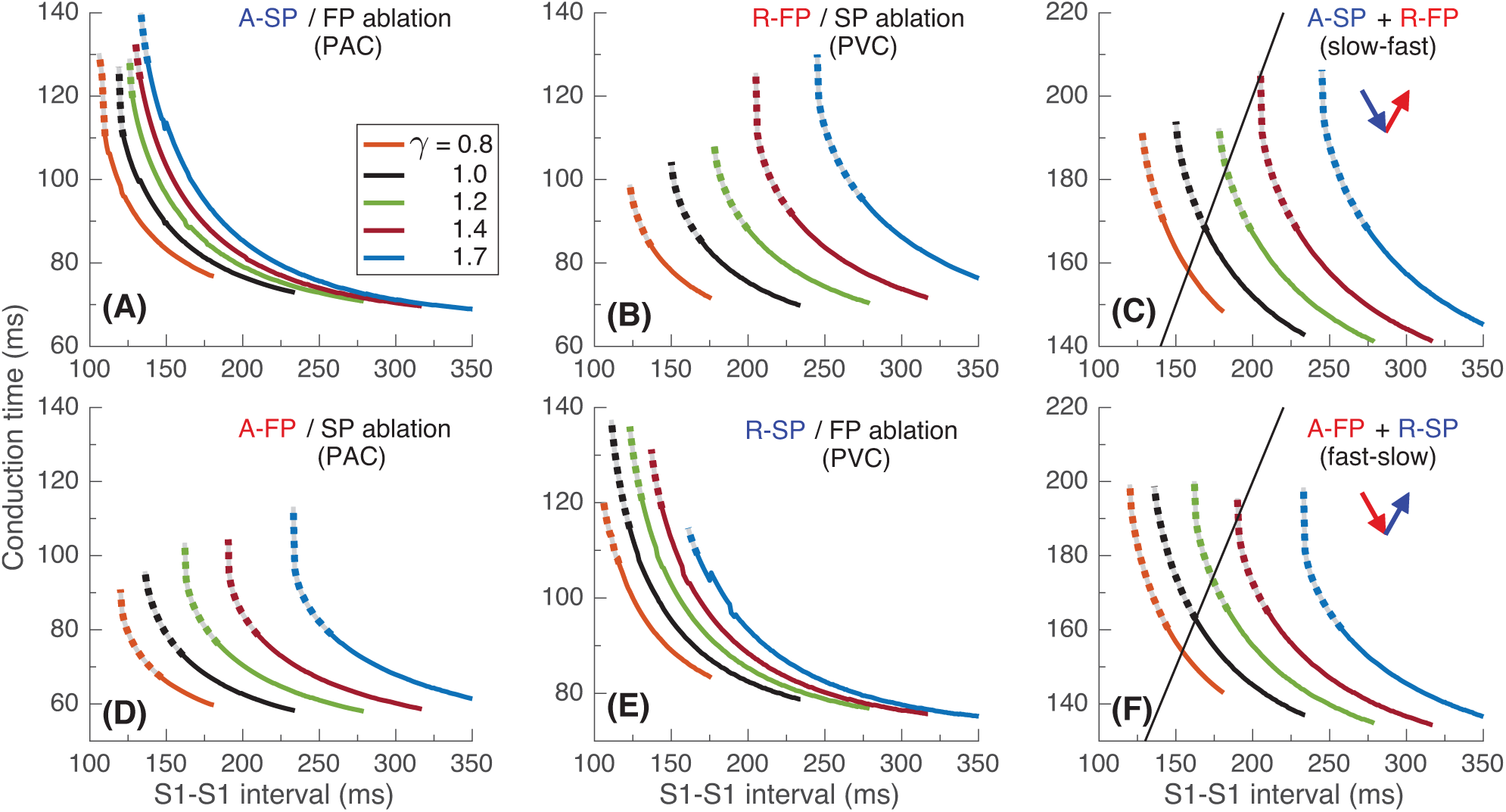
Conduction characteristics using S1S1 stimulation protocol for the second model variant.

### 2.3 The Third Model Variant

Figures 10–13 present simulation results obtained with the third model variant in which we set the relationship rERP*_SP_ >* rERP*_F P_* at enhanced sympathetic tone and its inversion (rERP*_SP_ <* rERP*_F P_*) at normal condition and enhanced vagal tone.

**Figure 10.**
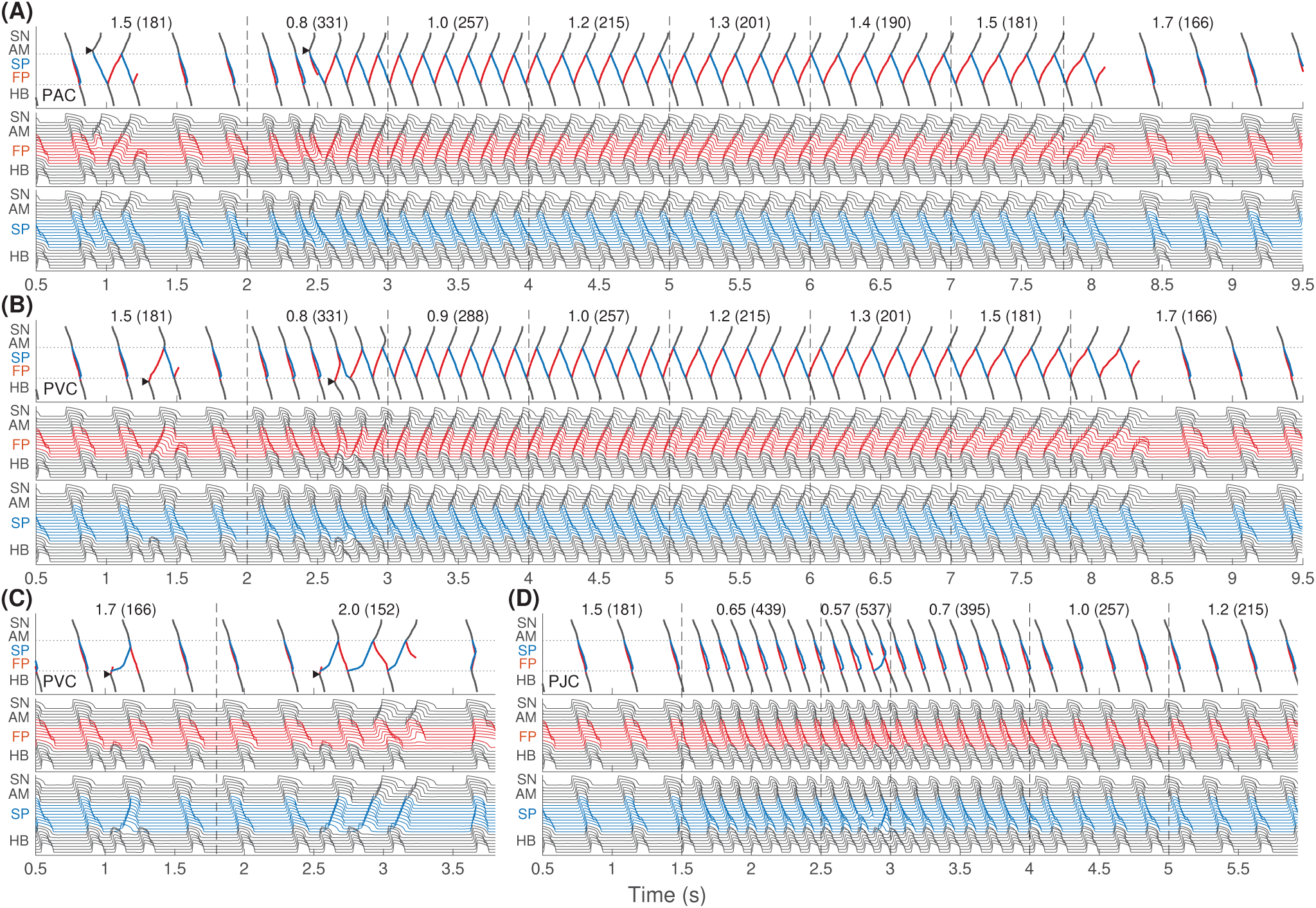
The same as in Fig. 2 but for the third model variant. Onset of slow-fast AVNRT with PAC (A) and with PVC (B) at enhanced sympathetic tone, and spontaneous termination of the oscillations at increasing vagal tone at *γ* ≥ 1.7. However, with atrial and His bundle stimulation at *γ* = 1.5 only fast-slow echo beats appeared. (C) Fast-slow echo beats also appeared with PVC at highly predominant parasympathetic tone (*γ* ≥ 1.7). (D) In contrast to Figs. 2 and 6, a brief burst of very strong sympathetic tone did not induce any AVNRT.

The situation with slow-fast AVNRT induction [Fig. 10(A)] looks the same as in Figs. 2(A) and 6(A) due to the similarity of anterograde conduction and refractory curves in panels (D)–(F) of Figs. 4, 8, and 12. However, with His bundle pacing [PVC, Fig. 10(B)] we obtained the same typical slow-fast AVNRT form as with atrial pacing [PAC, Fig. 10(A)]. At *γ* = 1.5, both slow-fast echo beat and AVNRT were initiated depending on preceding conditions. We also managed to induce some fast-slow echo beats at enhanced parasympathetic tone with *γ* = 1.7 − 2.0 [see Fig. 10(C)].

In contrast to the first and second model variants, with the third model variant no AVNRT was induced with brief burst of very strong sympathetic tone [Fig. 10(D)]. In Fig. 11 the details of the onset of slow-fast AVNRT with His bundle pacing (PVC) are shown on ladder diagrams.

**Figure 11.**
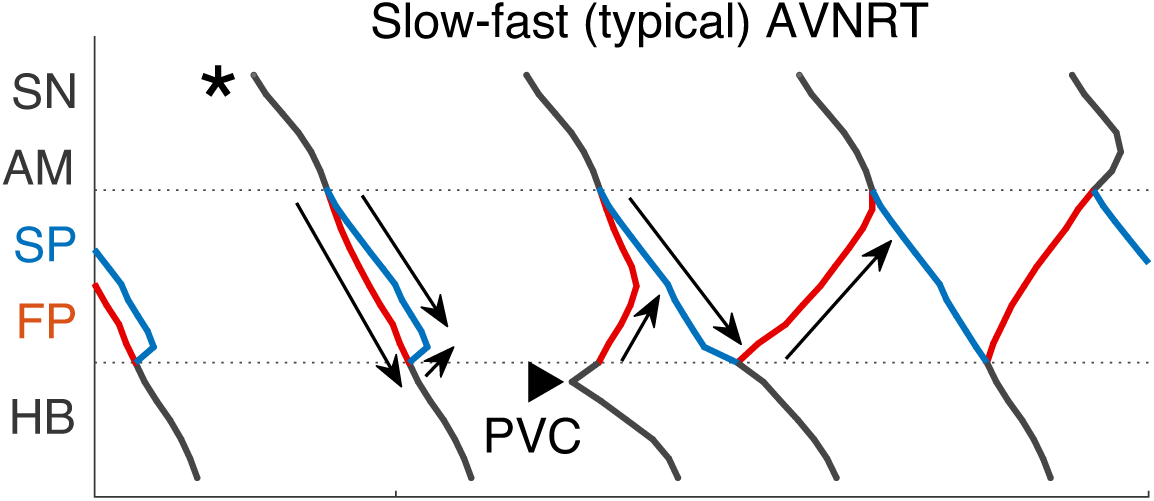
Scheme of slow-fast AVNRT onset with PVC in the third model variant [from Fig. 10(B)]. Arrows show the direction of conduction via pathways. Arrowhead denotes the place and moment of premature stimulation. Asterisk indicates normal conduction.

According to the setup of the third model variant, the retrograde FP and SP ERP curves shown in Fig. 12(I) intersect at *γ* ≃ 1.5, which suggests induction of slow-fast AVNRT in the *γ <* 1.5 range, and possible induction of fast-slow AVNRT at *γ >* 1.5. The peculiarity of the point *γ* = 1.5 is reflected in the unusual form of retrograde control and FP conduction curves in their leftmost points in Figs. 12(B) and 12(H).

**Figure 12.**
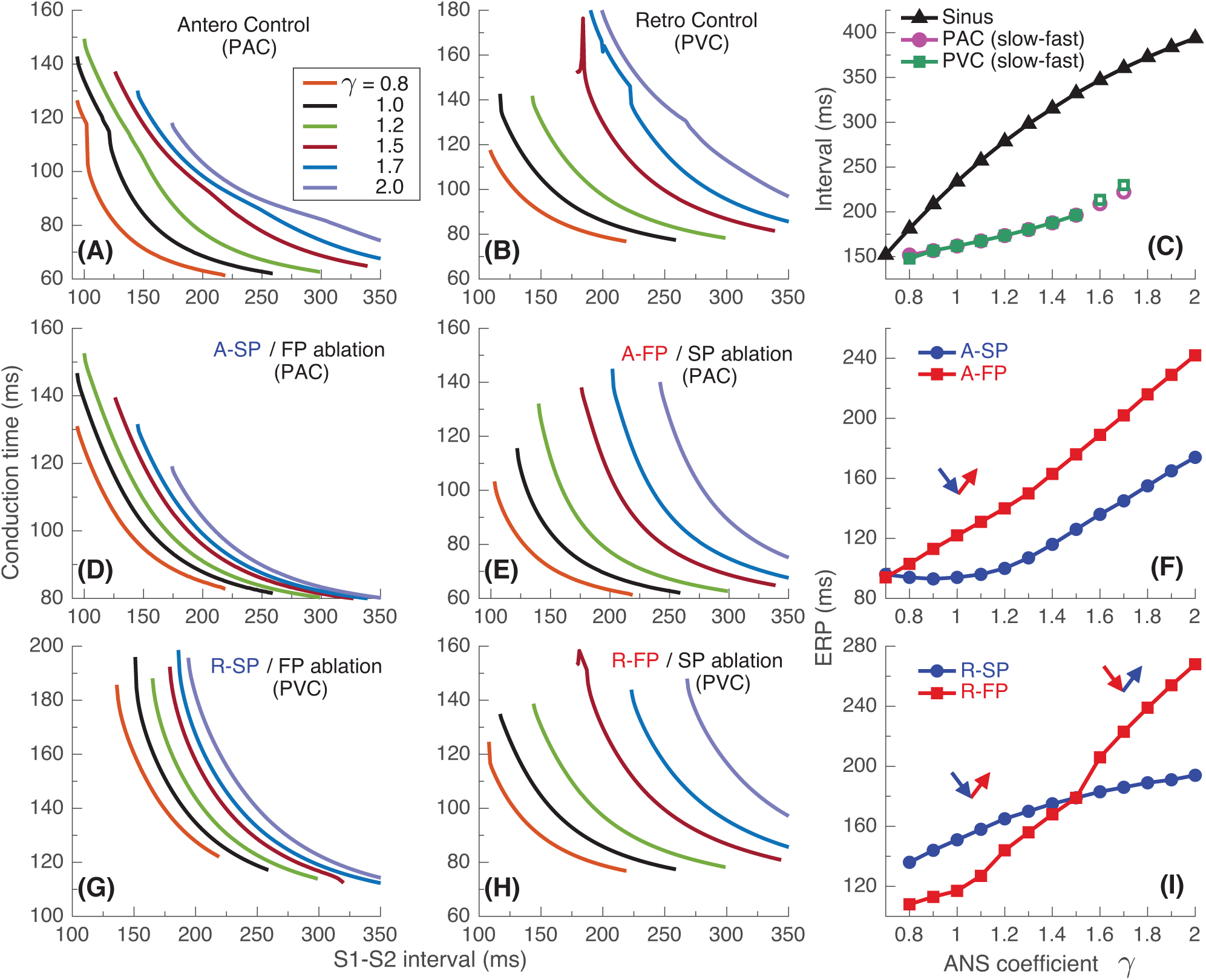
Conduction characteristics using S1S2 stimulation protocol for the third model variant.

Results using S1S1 stimulation are demonstrated in Fig. 13. Sustained slow-fast AVNRT can exist in the range *γ* ≤ 1.5 [Fig. 13(C)]. The fast-slow form can exist within the entire range of *γ* which is supported by intersections of the AVNRT sustainability line with the whole set of conduction curves [Fig. 13(F)], but it should be unstable at very high (1.7–2.0) values of *γ*.

**Figure 13.**
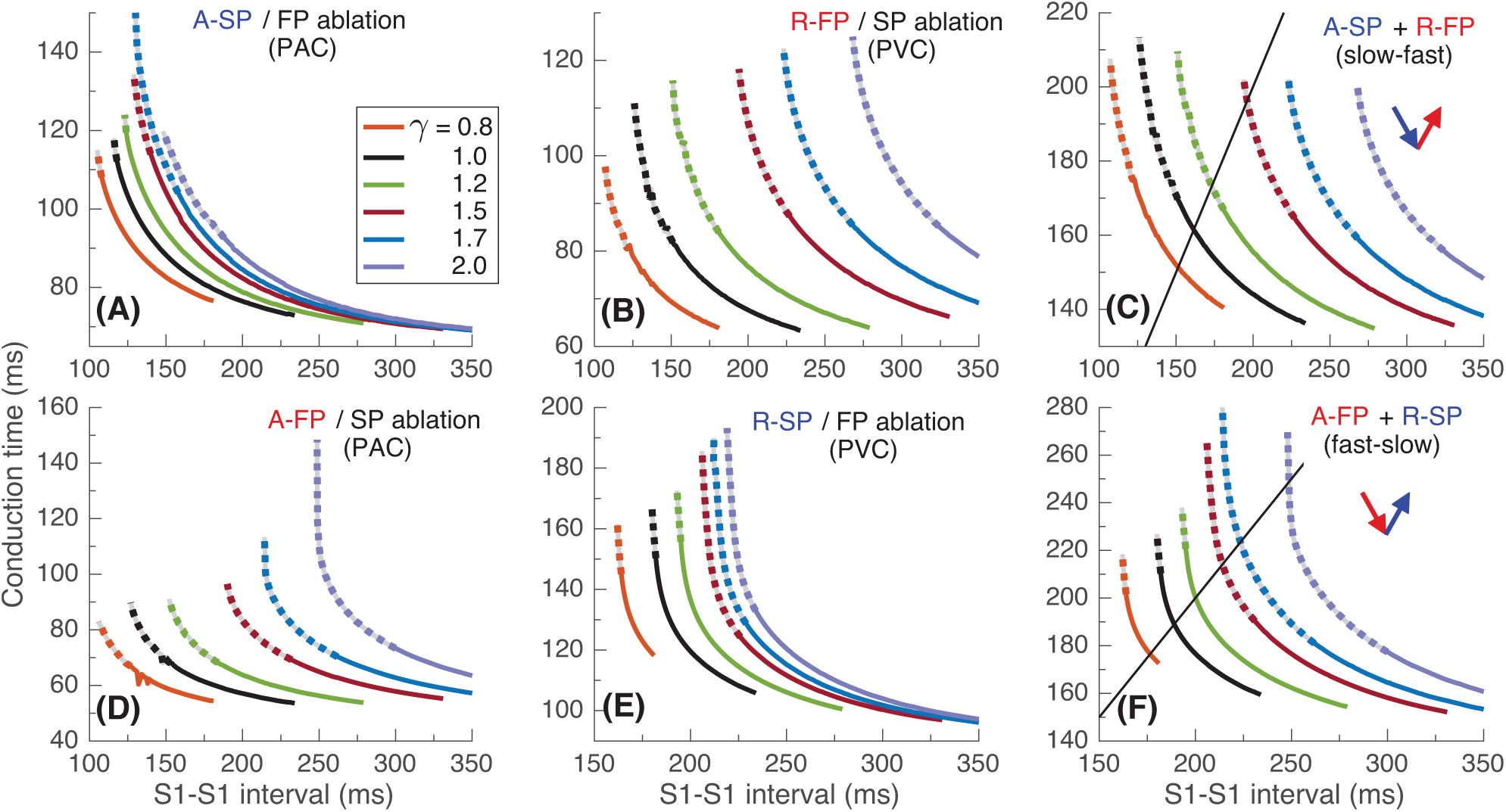
Conduction characteristics using S1S1 stimulation protocol for the third model variant.

## 3 DISCUSSION

Using our compact multifunctional model of rabbit AV node, we simulated the effect of ANS on the behavior of AVNRT. Incorporating a single ANS coefficient *γ* allowed the introduction of the combined effect of sympathetic and parasympathetic activity into the basic AV node model through the modulation of the refractoriness of model cells. A decrease in *γ* increases sympathetic tone, diminishes parasympathetic tone, and leads to a reduction in AV nodal conduction time and nodal refractory period in accordance with the results of electrophysiological studies (19, 8). On the other hand, parasympathetic activity dominates with increasing *γ* and has opposite effects on AV nodal conduction time and refractory period (18). The approach is somewhat similar to one used in the work (21). Still, our model utilizes a one-dimensional reduced-order reaction-diffusion system composed of 32 model cells and one ANS control coefficient.

The validity of our approach to the incorporation of ANS control into our rabbit AV node model and obtained results are supported by the following clinical and experimental observations.

Clinical studies suggest that sustained typical slow-fast AVNRT episodes are preceded by an increase in sympathetic tone (20). Accordingly, for the induction of AVNRT in most cases, we set *γ* to a small value (0.80–1.0, corresponding to enhanced sympathetic tone) and obtained slow-fast AVNRT [Figs. 2(A), 6(A), and 10(A)].

The atypical fast-slow AVNRT form appeared in our simulation in rare cases: in the case with PJC [Figs. 2(C) and 6(C)], occurring primarily in children and postoperative patients (4), in the case with PVC [Fig. 6(B)] with rERP*_SP_ <* rERP*_F P_* (31), and in even more marginal case with PVC [Fig. 10(C)] at high vagal tone (6). These results are supported by the clinical fact of a significant predominance of the typical slow-fast form of AVNRT than the atypical fast-slow one (17).

As described in (16), under certain conditions, the difference in retrograde ERPs between SP and FP may become inverted with the changing of the ANS state [Fig. 12(I)], resulting in both slow-fast and fast-slow AVNRT forms can present in the same subject. In Fig. 10(A), typical slow-fast AVNRT appeared at the predominance of sympathetic tone in *γ* ≤ 1.5 range. On the other hand, the atypical fast-slow echo beats were induced at enhanced vagal tone (*γ >* 1.7) [Fig. 10(C)]. The onset of AVNRT throughout periods of increased vagal tone, such as during sleep, was observed in clinical practice (6).

While the onset of AVNRT is mostly observed at enhanced sympathetic activity, a possibility exists for the tachycardia or reciprocating beats induction with increased vagal tone due to the widening of the induction window (6, 28). With increasing parameter *γ* and decreasing sinus rate, the inducibility of slow-fast AVNRT by atrial extrastimuli strengthened in all model variants [Figs. 4(F), 8(F), and 12(F)]. We also observed the same effect of ANS on the fast-slow AVNRT inducibility with His bundle stimulation for the second model variant [Fig. 8(I)].

The *onset* of AVNRT requires four factors: (A) existence of at least two functional AV nodal pathways, a specific ANS state providing proper refractoriness and conduction time of the AV nodal pathways, a difference in ERPs between the slow and fast pathways, and (D) premature atrial or His bundle (ventricular) stimulus with proper timing. The latter condition is unnecessary when AVNRT originated within the AV node (PJC). Induction of AVNRT by a brief burst of enhanced sympathetic tone accompanied by fast sinus rhythm or atrial pacing was reported in patients (6). This phenomenon was also observed in our simulations [Figs. 2(C) and 6(C)].

The *sustainability* of AVNRT is determined by the coincidence of the total duration of anterograde and retrograde conduction in slow and fast pathways with the stimulation period (AVNRT cycle length) at the certain state of the ANS [panels (C) and (F) in Figs. 5, 9, and 13]. If the AVNRT sustainability condition is not satisfied but the conditions for its onset mentioned in the previous paragraph are met, a few echo beats may occur [Figs. 6(B) and 8(I)].

As seen in Figs. 2(A), 6(B), and 10(a,b), at the same coefficient *γ* all kinds of behavior may be observed - a few echo beats, sustained tachycardia, or its termination, depending on the preceding activity. Thus, from the nonlinear dynamics point of view, the sustainability criteria appear to be a basin of attraction (14), when external stimuli at a particular set of initial conditions either lead to persistent oscillations within the AV node ring, which may be accompanied by cycle length variability (32), or to fading echo beats. The nonlinear analysis of the AVNRT behavior requires further study.

The shapes of anterograde control conduction curves obtained using S1S2 protocol with different coefficient *γ* can be with more or less marked tilting and bend at the point of conduction switching between FP and SP [Figs. 4(A), 8(A), and 12(A)] (22, 25). The bending and the nodal conduction discontinuity became more pronounced with smaller *γ*. The discontinuity position hardly observed at large *γ*, shifted toward a shorter S2 coupling interval due to a significant decrease of aERP*_F P_*. The smooth and discontinuous anterograde conduction curves were experimentally demonstrated in rabbits (22, 35) and in humans (31). The nodal conduction discontinuities appeared in some retrograde control conduction curves with PVC in the second model variant [Fig. 8(B)] and in the third model variant [Fig. 12(B)] were also observed in patients (33, 31) due to the prevalence of rERP*_F P_* over rERP*_SP_* [Figs. 8(I) and 12(I)].

In most cases of anterograde and retrograde conduction, the vagal modulation affected the ERPs of FP stronger than the ERPs of SP [see panels (F) and (I) in Figs. 4, 8, and 12], as quantitatively demonstrated in (6).

Applying S1S1 stimulation protocol, we observed some cases of nodal conduction alternans with the variation of conduction time from beat to beat (30, 9), which are reflected in some bumps on conduction curves in Figs. 5(A), 5(E), 9(A), 9(E), 13(B) and 13(D).

The *spontaneous termination* of AVNRT takes place with increasing refractoriness and nodal conduction delays (21) due to enhanced vagal tone, which is used in Valsalva meneuvers and pharmaceutical therapy (2, 34). When the sum of anterograde and retrograde conduction delays in SP and FP becomes not equal to the pacing interval, the condition for persistent AVNRT disappears [panels (C) and (F) in Figs. 5, 9, and 13]. In the vast majority of cases, spontaneous termination of AVNRT occurred when the conduction was blocked through FP regardless of the AVNRT form [Figs. 2(A), 2(C), 6, 10(A), 10(B), and 10(C)]. This is in line with clinical observations on the spontaneous AVNRT termination (5).

The summary of different types of AVNRT and echo beats obtained in our simulations is given in Table 1, where S-F and F-S mean slow-fast and fast slow types, and asterisk denotes echo beats. In the table, “Pulse” type corresponds to the onset of AVNRT induced by either PAC or PVC, and “ANS tone change” type is related to the sustainability of the oscillations. As seen from Table 1, at enhanced parasympathetic tone (*γ* ≥ 1.7) the reentrant activity decreases significantly and is represented only by echo beats.

**Table 1.**
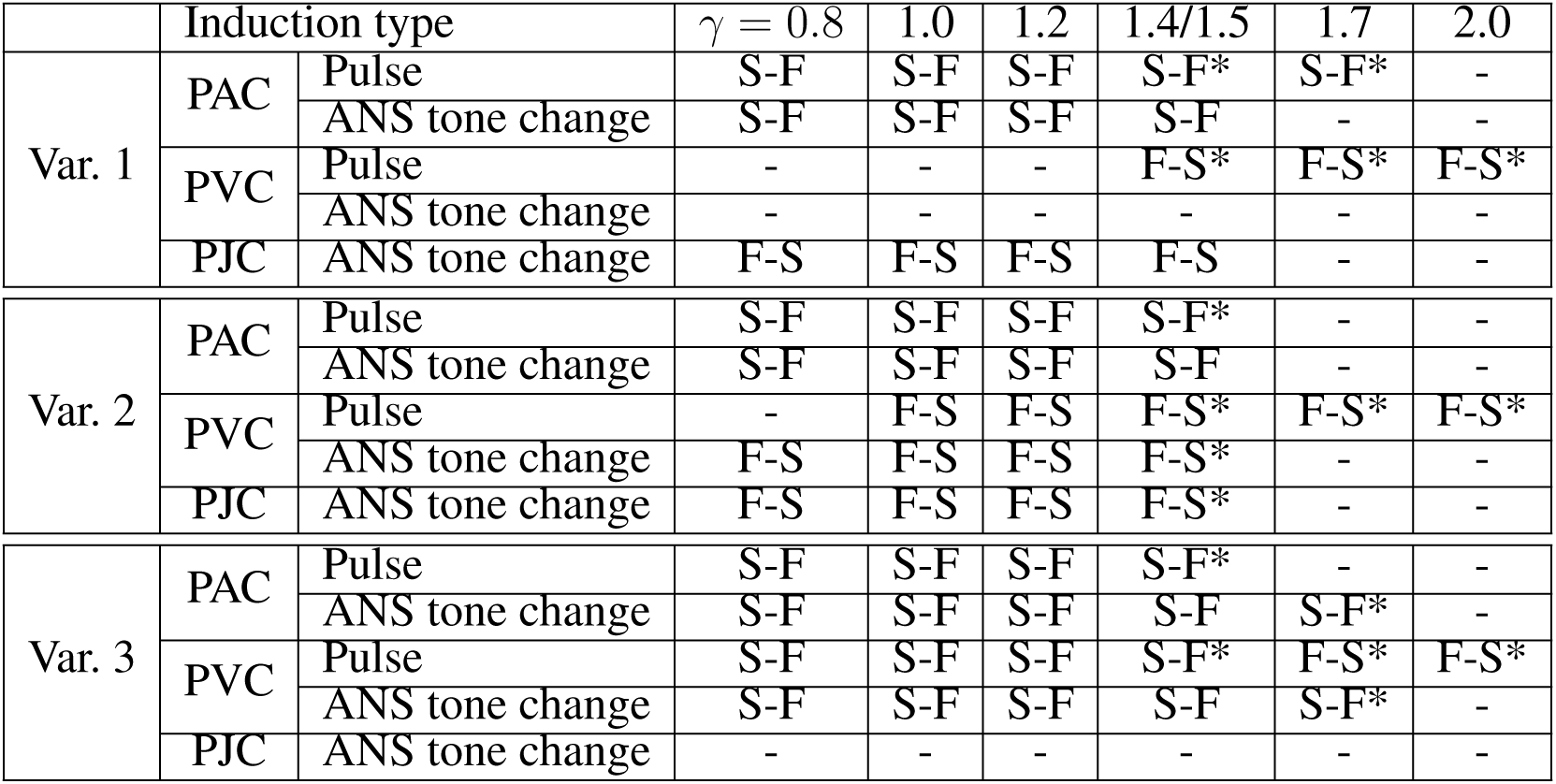
Induction of different types of AVNRT in the three model variants. Asterisk denotes echo beat(s).

| | Induction type | | $\gamma = 0.8$ | 1.0 | 1.2 | 1.4/1.5 | 1.7 | 2.0 |
| --- | --- | --- | --- | --- | --- | --- | --- | --- |
| Var. 1 | PAC | Pulse | S-F | S-F | S-F | S-F* | S-F* | - |
|  |  | ANS tone change | S-F | S-F | S-F | S-F | - | - |
|  | PVC | Pulse | - | - | - | F-S* | F-S* | F-S* |
|  |  | ANS tone change | - | - | - | - | - | - |
|  | PJC | ANS tone change | F-S | F-S | F-S | F-S | - | - |
| Var. 2 | PAC | Pulse | S-F | S-F | S-F | S-F* | - | - |
|  |  | ANS tone change | S-F | S-F | S-F | S-F | - | - |
|  | PVC | Pulse | - | F-S | F-S | F-S* | F-S* | F-S* |
|  |  | ANS tone change | F-S | F-S | F-S | F-S* | - | - |
|  | PJC | ANS tone change | F-S | F-S | F-S | F-S* | - | - |
| Var. 3 | PAC | Pulse | S-F | S-F | S-F | S-F* | - | - |
|  |  | ANS tone change | S-F | S-F | S-F | S-F | S-F* | - |
|  | PVC | Pulse | S-F | S-F | S-F | S-F* | F-S* | F-S* |
|  |  | ANS tone change | S-F | S-F | S-F | S-F | S-F* | - |
|  | PJC | ANS tone change | - | - | - | - | - | - |

One of the limitations of the current version of the rabbit conduction system model is the absence of heart rate variability. The heart rate variability is also controlled by the balance between parasympathetic and sympathetic tones of the ANS (11, 23). The second limitation is that we applied the ANS control coefficient *γ* to SP and FP on the same scale, which led to some differences in their response. In contrast, in reality, the degree of influence of ANS on the AV nodal pathways may differ. Finally, the current structure of the AV node model includes only one slow pathway. Thus, the atypical slow-slow form of AVNRT (17) was left out of our consideration.

## 4 CONCLUSION

In this work, we extended the functionality of our previously developed model of the rabbit cardiac conduction system based on the Aliev-Panfilov cardiac cell model by incorporating control from the autonomic nervous system. The control is accomplished by altering cell refractoriness using a single coefficient, which changes the conduction delay in the AV nodal pathways and intrinsic frequency of pacemaker cells. The influence of the autonomic nervous system extends from the sinoatrial node to the proximal part of the His bundle simultaneously in all cells of the model.

Using the modified model, we studied conditions for the onset, sustainability, and spontaneous termination of typical slow-fast and atypical fast-slow AVNRT forms. The conditions for the onset and sustainability of AVNRT can occur independently in various combinations. The difference in effective refractory periods between slow and fast pathways and the certain state of the autonomic nervous system determine the type of AVNRT and its sustainability with both atrial pacing and His bundle pacing.

The presented updated computationally lightweight but detailed model of rabbit cardiac conduction system with dual AV nodal pathways is suitable for studying physiological mechanisms of various forms of AVNRT. Inclusion of autonomic nervous system control into the model provides more lifelike functionality and allows realizing of various situations that are nearly impossible to reproduce in animal or human experiments. Our model could also serve as an educational tool to help students and practitioners visualize and understand the dynamic and complex interactions leading to AVNRT.

## DATA AVAILABILITY STATEMENT

The data that support the findings of this study are available from the corresponding author, MR, upon reasonable request.

## AUTHOR CONTRIBUTIONS

MR wrote the program code and performed computer simulations. ER analyzed the results. All authors contributed to the conception and design of the study and manuscript preparation; read and approved the submitted version.

## REFERENCES

1 Aliev, R. R. and Panfilov, A. V. (1996). A simple two-variable model of cardiac excitation. Chaos Solitons Fractals 7, 293–301. doi:10.1016/0960-0779(95)00089-5

2 Appelboam, A., Reuben, A., Mann, C., Gagg, J., Ewings, P., Barton, A., et al. (2015). Postural modification to the standard Valsalva manoeuvre for emergency treatment of supraventricular tachycardias (revert): a randomised controlled trial. Lancet 386, 1747–1753. doi:10.1016/S0140-6736(15)61485-4

3 Billette, J. and Tadros, R. (2019). An integrated overview of AV node physiology. Pacing Clin. Electrophysiol. 42, 805–820. doi:10.1111/pace.13734

4 Chen, H., Shehata, M., Cingolani, E., Chugh, S. S., Chen, M., and Wang, X. (2015). Differentiating atrioventricular nodal re-entrant tachycardia from junctional tachycardia: conflicting responses? Circ. Arrhythm. Electrophysiol. 8, 232–235. doi:10.1161/CIRCEP.114.002169

5 Chiale, P. A., Baranchuk, A., González, M. D., Sánchez, R. A., Garro, H. A., Fernández, P. A., et al. (2015). The mechanisms of spontaneous termination of reentrant supraventricular tachycardias. Int. J. Cardiol. 191, 151–158. doi:10.1016/j.ijcard.2015.04.239

6 Chiou, C.-W., Chen, S.-A., Kung, M.-H., Chang, M.-S., and Prystowsky, E. N. (2003). Effects of continuous enhanced vagal tone on dual atrioventricular node and accessory pathways. Circulation 107, 2583–2588. doi:10.1161/01.CIR.0000068339.04731.4D

7 Climent, A. M., Guillem, M. S., Zhang, Y., Millet, J., and Mazgalev, T. N. (2011). Functional mathematical model of dual pathway AV nodal conduction. Am. J. Physiol. Heart Circ. Physiol. 300, H1393–H1401. doi:10.1152/ajpheart.01175.2010

8 Cossú, S. F., Rothman, S. A., Chmielewski, I. L., Hsia, H. H., Vogel, R. L., Miller, J. M., et al. (1997). The effects of isoproterenol on the cardiac conduction system: site-specific dose dependence. J. Cardiovasc. Electrophysiol. 8, 847–853. doi:10.1111/j.1540-8167.1997.tb00845.x

9 Garfinkel, A. (2007). Eight (or more) kinds of alternans. J. Electrocardiol. 40, S70–S74. doi:10.1016/j.jelectrocard.2007.06.011

10 George, S. A., Faye, N. R., Murillo-Berlioz, A., Lee, K. B., Trachiotis, G. D., and Efimov, I. R. (2017). At the atrioventricular crossroads: dual pathway electrophysiology in the atrioventricular node and its underlying heterogeneities. Arrhythmia Electrophysiol. Rev. 6, 179–185. doi:10.15420/aer.2017.30.1

11 Guzzetti, S., Borroni, E., Garbelli, P. E., Ceriani, E., Bella, P. D., N, N. M., et al. (2005). Symbolic dynamics of heart rate variability: A probe to investigate cardiac autonomic modulation. Circulation 112, 465–470. doi:10.1161/CIRCULATIONAHA.104.518449

12 Hartikainen, J. E., Kautzner, J., Malik, M., and Camm, A. J. (1997). Sympathetic predominance of cardiac autonomic regulation in patients with left free wall accessory pathway and orthodromic atrioventricular reentrant tachycardia. Eur. Heart J. 18, 1966–1972. doi:10.1093/oxfordjournals.eurheartj.a015207

13 Inada, S., Hancox, J. C., Zhang, H., and Boyett, M. R. (2009). One-dimensional mathematical model of the atrioventricular node including atrio-nodal, nodal, and nodal-His cells. Biophys. J. 97, 2117–2127. doi:10.1016/j.bpj.2009.06.056

14 Izhikevich, E. M. (2006). Dynamical systems in neuroscience: The geometry of excitability and bursting (The MIT Press). doi:10.7551/mitpress/2526.001.0001

15 Katritsis, D. G. and Josephson, M. E. (2013). Classification of electrophysiological types of atrioventricular nodal re-entrant tachycardia: A reappraisal. EP Europace 15, 1231–1240. doi:10.1093/europace/eut100

16 Katritsis, D. G., Marine, J. E., Latchamsetty, R., Zografos, T., Tanawuttiwat, T., Sheldon, S. H., et al. (2015). Coexistent types of atrioventricular nodal re-entrant tachycardia implications for the tachycardia circuit. Circ. Arrhythm. Electrophysiol. 8, 1189–1193. doi:10.1161/CIRCEP.115.002971

17 Katritsis, D. G., Sepahpour, A., Marine, J. E., Katritsis, G. D., Tanawuttiwat, T., Calkins, H., et al. (2015). Atypical atrioventricular nodal reentrant tachycardia: prevalence, electrophysiologic characteristics, and tachycardia circuit. Europace 17, 1099–1106. doi:10.1093/europace/euu387

18 Martin, P. (1977). The influence of the parasympathetic nervous system on atrioventricular conduction. Circ. Res. 41, 593–599. doi:10.1161/01.res.41.5.593

19 Morady, F., Nelson, S. D., Kou, W. H., Pratley, R., Schmaltz, S., Buitleir, M. D., et al. (1988). Electrophysiologic effects of epinephrine in humans. J. Am. Coll. Cardiol. 11, 1235–1244. doi:10.1016/0735-1097(88)90287-2

20 Nigro, G., Russo, V., de Chiara, A., A, A. R., Cioppa, N. D., Chianese, R., et al. (2010). Autonomic nervous system modulation before the onset of sustained atrioventricular nodal reentry tachycardia. Ann. Noninvasive Electrocardiol. 15, 49–55. doi:10.1111/j.1542-474X.2009.00339.x

21 Plappert, F., Wallman, M., Abdollahpur, M., Platonov, P. G., Östenson, S., and Sandberg, F. (2022). An atrioventricular node model incorporating autonomic tone. Front. Physiol. 13, 976468. doi:10.3389/fphys.2022.976468

22 Reid, M. C., Billette, J., Khalife, K., and Tadros, R. (2003). Role of compact node and posterior extension in direction-dependent changes in atrioventricular nodal function in rabbit. J. Cardiovasc. Electrophysiol. 14, 1342–1350. doi:10.1046/j.1540-8167.2003.03382.x

23 Rovere, M. T. L., Porta, A., and Schwartz, P. J. (2020). Autonomic control of the heart and its clinical impact. a personal perspective. Front. Physiol. 11, 582. doi:10.3389/fphys.2020.00582

24 Ryzhii, M. and Ryzhii, E. (2022). Pacemaking function of two simplified cell models. PLoS ONE 17, e0257935. doi:10.1371/journal.pone.0257935

25 Ryzhii, M. and Ryzhii, E. (2023). A compact multi-functional model of the rabbit atrioventricular node with dual pathways. Front. Physiol. 14, 1126648. doi:10.3389/fphys.2023.1126648

26 Ryzhii, M. and Ryzhii, E. (2023). Revealing the origin of typical and atypical forms of atrioventricular nodal reentrant tachycardia with a compact computer model of rabbit AV node. In Computing in Cardiology (CinC2023) (IEEE), 1–4. doi:10.22489/CinC.2023.080

27 Ryzhii, M. and Ryzhii, E. (2024). Compact computer model of rabbit atrioventricular node with autonomic nervous system control. In Computing in Cardiology (CinC2024) (IEEE), 1–4

28 Sinkovec, M., Pernat, A., Rajković, Z., Jan, M., Antolic, B., and Rakovec, P. (2011). Electrophy- siology of anterograde right-atrial and left-atrial inputs to the atrioventricular node in patients with atrioventricular nodal re-entrant tachycardia. Europace 13, 869–875. doi:10.1093/europace/euq459

29 Straus, D. G. and Schocken, D. D. (2021). Marriott’s Practical Electrocardiography, 13th *ed*. (Philadelphia: Wolters Kluwer Health)

30 Sun, J., Amellal, F., Glass, L., and Billette, J. (1995). Alternans and period-doubling bifurcations in atrioventricular nodal conduction. J. Theor. Biol. 173, 79–91. doi:10.1006/jtbi.1995.0045

31 Sung, R. J., Styperek, J. L., Myerburg, R. J., and Castellanos, A. (1978). Initiation of two distinct forms of atrioventricular nodal reentrant tachycardia during programmed ventricular stimulation in man. J. Am. Coll. Cardiol. 42, 404–415. doi:10.1016/0002-9149(78)90935-9

32 Tamura, S., Nakajima, T., Iizuka, T., Hasegawa, H., Kobari, T., Kurabayashi, M., et al. (2020). Unique electrophysiological properties of fast–slow atrioventricular nodal reentrant tachycardias characterized by a shortening of retrograde conduction time via a slow pathway manifested during atrial induction. J. Cardiovasc. Electrophysiol. 31, 1420–1429. doi:10.1111/jce.14501

33 Wu, D., ad F. Amat-Y-Leon, P. D., Wyndham, C. R., Dhingra, R., and Rosen, K. M. (1977). An unusual variety of atrioventricular nodal re-entry due to retrograde dual atrioventricular nodal pathways. Circulation 56, 50–59. doi:10.1161/01.cir.56.1.50

34 Xiao, L., Ou, X., Liu, W., Lin, X., Lin, P., Qiu, S., et al. (2024). Combined modified Valsalva maneuver with adenosine supraventricular tachycardia: A comparative study. Am. J. Emerg. Med. 78, 157–162. doi:10.1016/j.ajem.2024.01.035

35 Zhang, Y. (2016). His electrogram alternans (zhang’s phenomenon) and a new model of dual pathway atrioventricular node conduction. J. Interv. Card. Electrophysiol. 45, 19–28. doi:10.1007/s10840-015-0079-0

